# Protective microbe enhances colonisation of a novel host species by modifying immune gene expression

**DOI:** 10.1101/857540

**Authors:** Suzanne A. Ford, Kayla C. King

## Abstract

Microbes that protect against infection inhabit hosts across the tree of life. It is unclear whether many protective microbes use or reduce the need for a host immune response, or how the immune system reacts when these microbes newly encounter a host species naturally and as part of a biocontrol strategy. We sequenced the transcriptome of a host (*Caenorhabditis elegans*) following its interaction with a non-native bacterium (*Enterococcus faecalis*) that has protective traits against the pathogen, *Staphylococcus aureus*. We show that microbe-mediated protection caused the differential expression of 1,557 genes, including the upregulation of many immune gene families conserved across the animal kingdom (e.g. lysozymes and c-type lectins). We found that this modulation of the host’s immune response was beneficial for both the protective microbe and the host. Given *E. faecalis*’ increased ability to resist lysozyme activity compared to *S. aureus*, our results indicate that the protective microbe could more easily invade and protect infected hosts by upregulating lysozyme genes. These results suggest that a protective microbe can exploit the host immune system even when introduced into a novel species. Microbes that protect via the host immune response in this way should favour continued investment into host immunity and avoid the evolution of host dependence.

**Author summary:** Organisms can be protected from infectious disease by the microbes they house. It is unclear, however, whether protective microbes affect the host immune response to infection, particularly in the early stages of symbiosis. In this study, we investigated the role of the host immune system in a novel protective interaction. We examined gene expression in a nematode after colonisation by a non-native microbe capable of suppressing the pathogen *Staphylococcus aureus*. The protective microbe altered the host immune response to infection in a way that it could exploit. By causing the host to increase the production of antimicrobials to which it itself is relatively resistant, the protective microbe was better able to colonise and defend infected hosts. These results indicate that protective microbes introduced into new host species can take advantage of the host immune system. Such a mechanism at the beginning of a protective symbiosis, formed either naturally or as part of a biocontrol strategy, could ensure continued investment in host-based defences over evolutionary time.

## Introduction

Microbes that defend against infection by pathogens (along with parasites and parasitoids) inhabit a large diversity of plant and animal hosts(1, 2). Protection is most commonly found when symbiotic microbes suppress pathogen growth and establishment by directly competing for resources/space or producing antimicrobial compounds/toxins(2, 3). Microbes can also control infection by modulating host immune systems(2, 3). Modification of the host’s immune response against infection is a less commonly documented protective mechanism(2), although is unclear whether this pattern is real as immune-mediation may be more difficult to empirically test in non-model systems. An example is seen in the tropical tree *Theobroma cacao* where the foliar endophytic fungus *Colletotrichum tropicale* upregulates host immune and defence genes increasing resistance to damage from pathogens(4).

Interest in the mechanisms underpinning microbe-mediated protection has been surging because of its potential use in public health(5), species conservation(6) and agriculture(7). Although new protective symbioses form naturally(8–11), their creation is being rapidly pursued for the biocontrol of infectious disease, either by introducing existing symbionts into new hosts(12), or by generating new symbionts, e.g. via paratransgenesis(13–15). For example, *Aedes* and *Anopheles* mosquitoes have been artificially infected with strains of the inherited symbiont *Wolbachia* that inhibit the replication of dengue and Zika viruses, as well as malaria(5, 16–18). In addition to determining how a microbe might protect a novel host, the specific mechanism of protection might predict the persistence of the microbial symbiont(2), as well as the vulnerability of the host to infection should the mutualism break down(2, 19–22). If protective microbes directly suppress infection, there is less need for hosts to defend themselves. This outcome could result in reduced investment into costly host-based immune or damage response systems(1, 2, 23, 24). Conversely, a mechanism of protection that involves the immune response is likely to favour continued investment by the host(4). It is thus important to assess at the origin of a novel protective symbiosis whether the immune response is useful in protection or made redundant.

In this study, we disentangled the impacts of protective microbes on the immune system of a novel host species. We measured whole-genome transcriptional changes in the model host *Caenorhabditis elegans* upon colonisation by a non-native bacterium (*Enterococcus faecalis*) that has protective traits in other animals (25, 26). Because this host system is genetically tractable with a suite of available immune knock-out mutants, it is useful for exploring the role of the immune response amidst microbe-mediated protection. We have previously shown that a human-derived isolate of *E. faecalis* can colonise *C. elegans* nematodes and increase host survival during virulent infection by the opportunistic pathogen *Staphylococcus aureus* (27). Although we have demonstrated that this protective bacterium can quickly evolve to produce antimicrobial reactive oxygen species at levels inhibitory to *S. aureus* (27), other mechanisms are likely at play at the first point of interaction. During co-colonisation, *E. faecalis* can inhibit *S. aureus* growth by stealing iron-binding siderophores that the pathogen releases into the environment (28), and when colonising on its own, *E. faecalis* can upregulate a variety of immune genes in *C. elegans* (29).

We exposed *C. elegans* populations to the pathogen with and without *E. faecalis*-mediated protection, and we then measured gene expression profiles after 12 hours using RNA-sequencing technology (Fig.1). We found that protection drove the upregulation of numerous key *C. elegans* immune genes conserved across the animal kingdom, including 20 c-type lectin genes and six lysozyme genes. By disproportionately targeting the pathogen, expression of the lysozyme gene *lys-7* gave a competitive advantage to *E. faecalis* in infected hosts, thereby enhancing its protective ability. That this protective microbe was able to exploit the immune system of a novel host species suggests the potential for the immune system to be involved in the formation of protective symbioses more widely in nature or as part of a biocontrol strategy.

**Figure 1.**
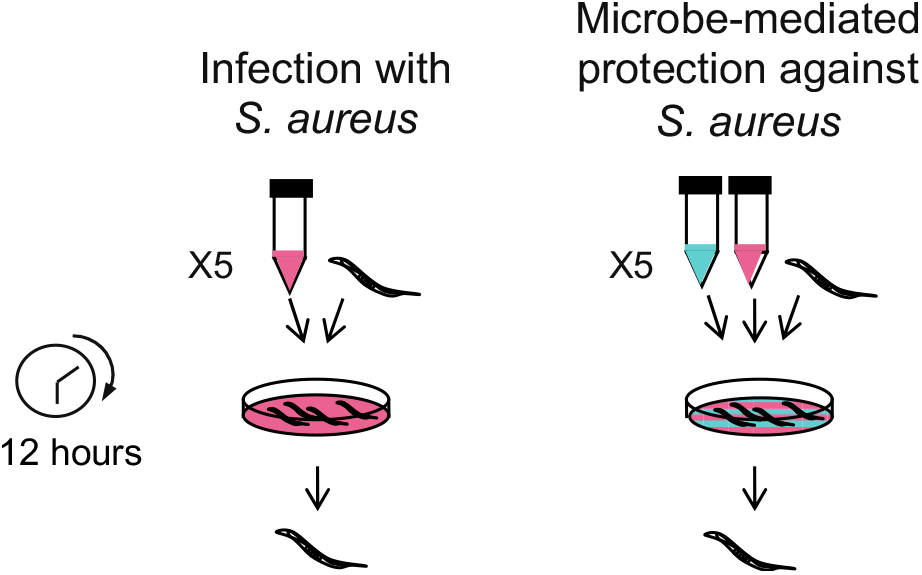
Experimental design for testing the effect of a novel protective microbe on the *C. elegans’* response to infection. Populations of young adult worms were exposed to the pathogen *S. aureus* with or without *E. faecalis-mediated* protection. After 12h exposure, the RNA from approximately 1,000 worms per sample was sequenced. This was repeated five independent times for each treatment.

## Results

### *E. faecalis*-mediated protection shapes *C. elegans’* response to infection

We used RNA-sequencing to assess the transcriptional changes in *C. elegans* hosts during infection by virulent *S. aureus*, with and without protection from *E. faecalis* (Fig.1). We found that protection drove the significant differential expression of 1,557 genes (Supplementary File 1). Of these genes, 521 were downregulated in protected hosts, whilst 1,036 were upregulated. We performed a GO-term enrichment analysis on this list of genes and identified significantly enriched GO-terms in biological processes including “defence response to bacterium” (GO:0042742, FDR-adjusted P= 6.12E-09, Supplementary File 2) and “innate immune response” (GO:0045087, FDR-adjusted P= 4.09E-12, Supplementary File 2). These are illustrated in Figure 2. We also found significantly enriched GO-terms in molecular functions including ‘structural constituent of cuticle’ (FDR-adjusted P=1.02E-37, Supplementary File 2).

**Figure 2.**
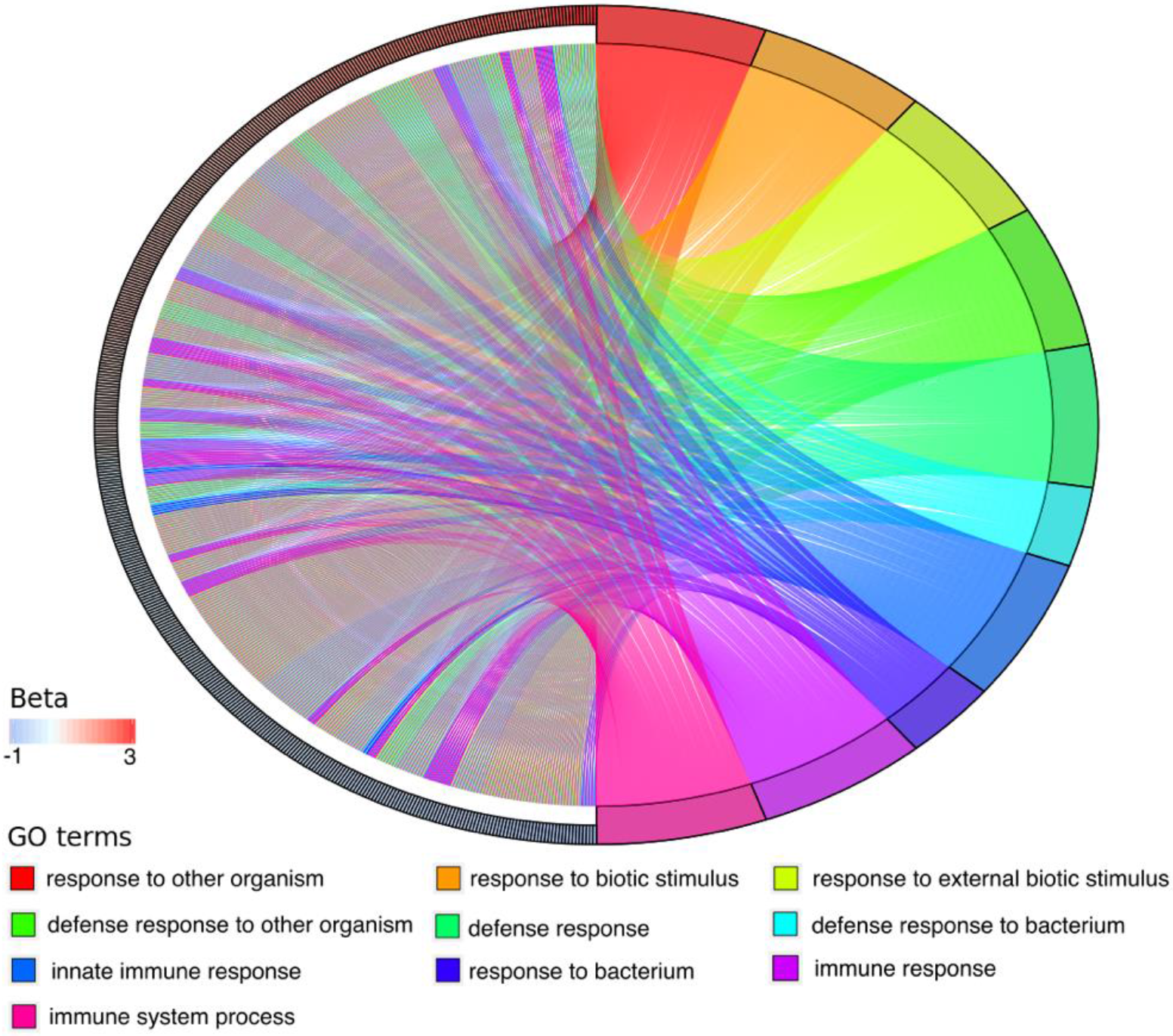
Significantly enriched gene ontology (GO)-terms for genes differentially expressed in protected hosts. A chord plot where *C. elegans* differentially expressed genes (left of circle) are linked by chords to the GO-terms (right of circle). The differentially expressed genes are ordered by beta values. Positive beta values indicate here the effect size of upregulation of a gene caused by microbe-mediated protection, whilst negative values indicate downregulation.

### Novel protective microbe upregulates immune genes in infected hosts

We identified many lysozyme-encoding genes that were differentially expressed during infection in *E. faecalis-protected* hosts (Table 1). Lysozymes are important in the host defence response to bacteria as these enzymes break down bacterial cell walls(30). We found that protection by *E. faecalis* caused the upregulation of the invertebrate lysozyme genes, *ilys-2* and *ilys-5* compared to when worms were only infected by *S. aureus (ilys-2* beta=1.171, P=6.28E-06; *ilys-5* beta=0.85, P=0.00055). Previous studies found that RNAi knockdown of *ilys-2* made *C. elegans* hosts more susceptible to *S. aureus* infection(31).

**Table 1.**
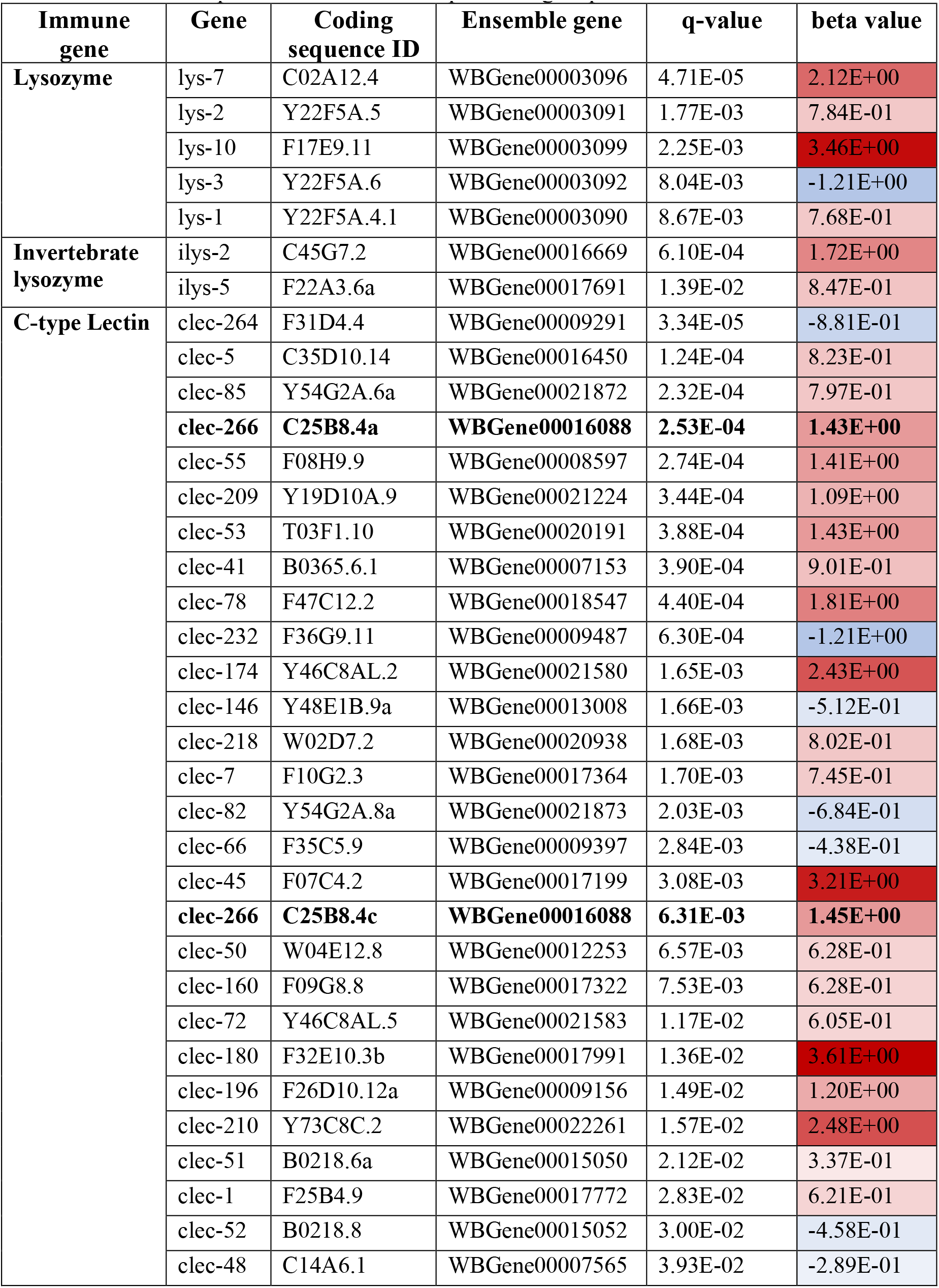
Immune gene families differentially regulated by *E. faecalis*-mediated protection in *C. elegans*. Genes are listed in order of q-value (p-values corrected by Benjamini-Hochberg FDR for multiple testing). The beta value is the effect size of differential regulation. Blue indicates downregulation whilst red indicates upregulation. Genes in bold are represented twice as unique coding sequences.

We also found a significant upregulation in the lysozyme genes, *lys-1* (beta=0.77, P=0.00028), *lys-2* (beta=0.78, P=3.01E-05), *lys-7* (beta=2.12, P=1.15E-07) and *lys-10* (beta=3.46, P=4.36E-05). Only *lys-3* was significantly downregulated (beta=−1.2, P=0.00025). The most significantly upregulated lysozyme gene was *lys-7*. Although both *E. faecalis* and *S. aureus* have been reported to be resistant to lysozyme activity, *E. faecalis* is much more robust to attack by these enzymes, with a minimum inhibitory concentration (MIC) of >62.5mg/ml lysozyme(32). *S. aureus* has lower reported MICs of 15mg/ml(33).

Microbe-mediated protection also resulted in the differential regulation of 28 C-type lectin (*clec*) genes which are carbohydrate-binding proteins that play a host defensive role against gram-positive bacteria(34). We found that 20 *clec* genes were upregulated, whilst only seven were downregulated (Table 1). Infection by *S. aureus* has been previously shown to upregulate *clec* genes in *C. elegans*, e.g. *clec-72* and *clec-52* (35). In our study, *clec-72* was upregulated in protected hosts, whilst *clec-52* was downregulated.

We additionally found numerous *C. elegans* cuticle genes that were differentially regulated by protection. These genes were entirely upregulated and consisted of 44 *col* and 8 *dpy* genes (Supplementary File 1). Both *col* and *dpy* genes are involved in the extracellular matrix and the cuticle. It is unclear what role these genes play in response to microbe-mediated protection, but genes encoding structural constituents of the cuticle are found to be significantly downregulated during *S. aureus* infection alone (35).

### Lysozyme expression determines protective microbe colonisation in novel infected host

Lysozymes are a well-known conserved antimicrobial defence(30). Given that *E. faecalis* is reported to be more resistant to lysozyme activity than *S. aureus*(32, 33), we tested whether the upregulation of lysozyme genes by *E. faecalis* played a role in protection against *S. aureus*. We focused on the impact of the most significantly differentially expressed lysozyme gene, *lys-7*, on bacterial colonisation and protection.

We obtained a *lys-7* gene knock-out in an N2 background (strain CB6738, *Caenorhabditis* Genetics Center, Minnesota) and tested whether *lys-7* affected host mortality and bacterial colonisation during microbe-mediated protection from infection. We found that host mortality was higher in *lys-7* knock-out hosts than the wild-type during co-colonisation with *E. faecalis* and *S. aureus*, indicating that it plays a role in resistance amidst microbe-mediated protection (Fig 3a. Quasibinomial GLM. host strain: F=14.7, df=1, P= 0.005). We found that *E. faecalis* was significantly better at colonising wild-type hosts than the *lys-7* knock-out mutant (Fig 3b. t-test, t=−3.5, df=8, P= 0.0076). By contrast, *S. aureus* reached higher infection loads in the *lys-7* knock-out mutant (Fig 3b. t-test, t=1.42, df=8, P=0.19). These data indicate that expression of *lys-7* yields a host-derived antimicrobial that negatively effects *S. aureus* more than *E. faecalis* under co-colonisation. This differential effect is consistent with the finding that *E. faecalis* has a much higher MIC for lysozyme than *S. aureus*(32, 33), and this may have enhanced *E. faecalis’* ability to colonise and compete within infected hosts.

**Figure 3.**
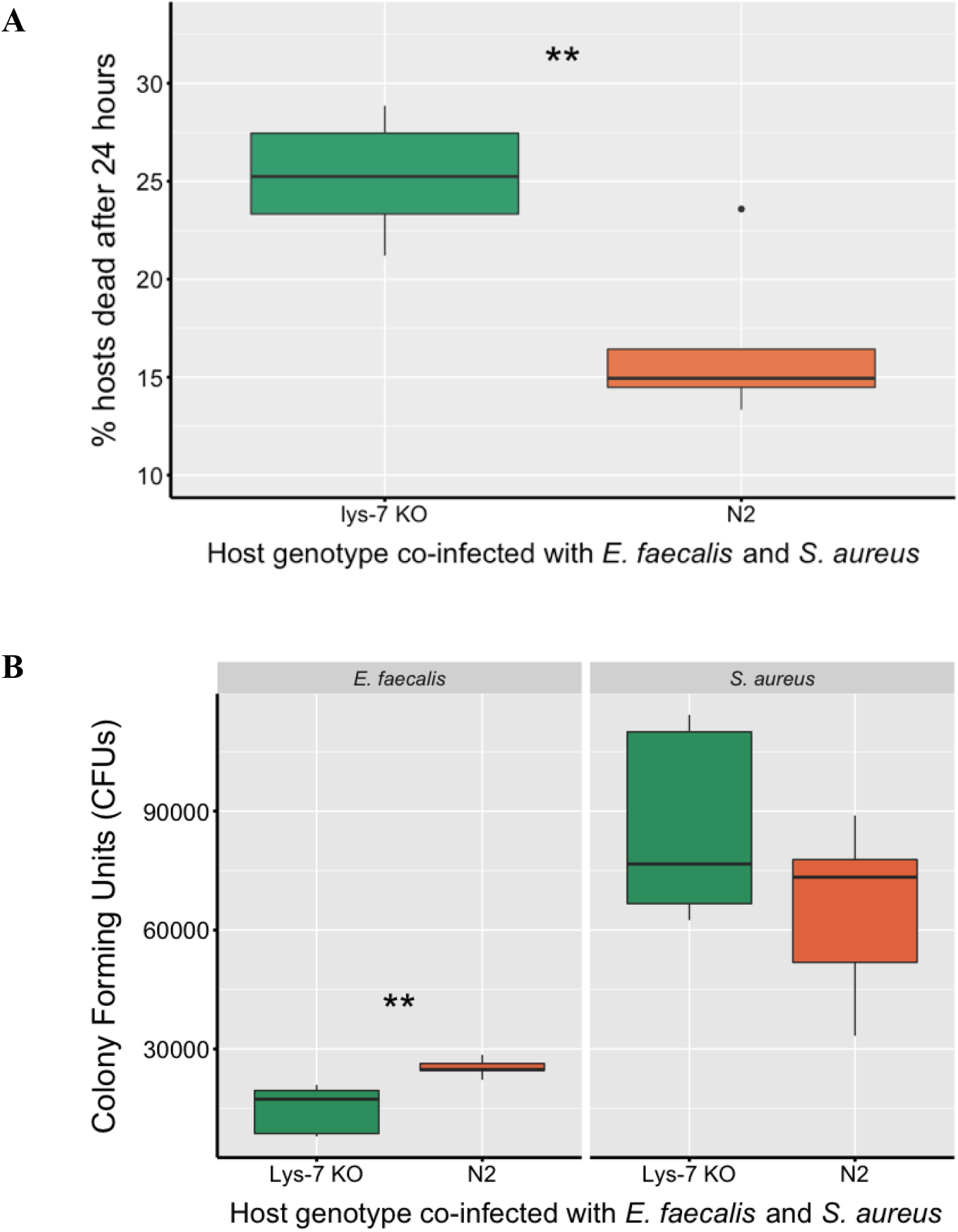
The effect of the *lys-7* gene on host mortality and bacterial colonisation during microbe-mediated protection against *S. aureus*. **A**, The percentage mortality of wild-type N2 *C. elegans* hosts and the *lys-7* knock-out mutant after 24 hours of co-colonisation by *S. aureus* and *E. faecalis*. Each treatment was replicated five independent times. **B**, Within-host bacterial load in colony forming units (CFUs) of *E. faecalis* and *S. aureus* after 24 hours of exposure in the wild-type N2 host and *lys-7* knockout mutant. Each treatment was replicated five independent times and 7-10 worms were collected per replicate. *P<0.05, **P<0.01, ***P<0.001.

To test whether the lysozymes produced by *lys-7* expression could affect host resistance to *E. faecalis* and *S. aureus*, we measured the mortality of the wild-type N2 and *lys-7* knockout mutant when exposed to each bacterium. We found that *lys-7* played a significant role in host resistance to all bacterial exposure treatments (including to food), however we saw the largest change in host mortality in response to *S. aureus* infection to almost 100% of the nematode population after 24h (Fig 4. control OP50 food: Binomial GLM, df=1, FDR-corrected P=0.04; *E. faecalis:* Binomial GLM, df=1, FDR-corrected P=0.04; *S. aureus:* Quasibinomial GLM, F=11.2, df=1, FDR-corrected P= 0.03).

**Figure 4.**
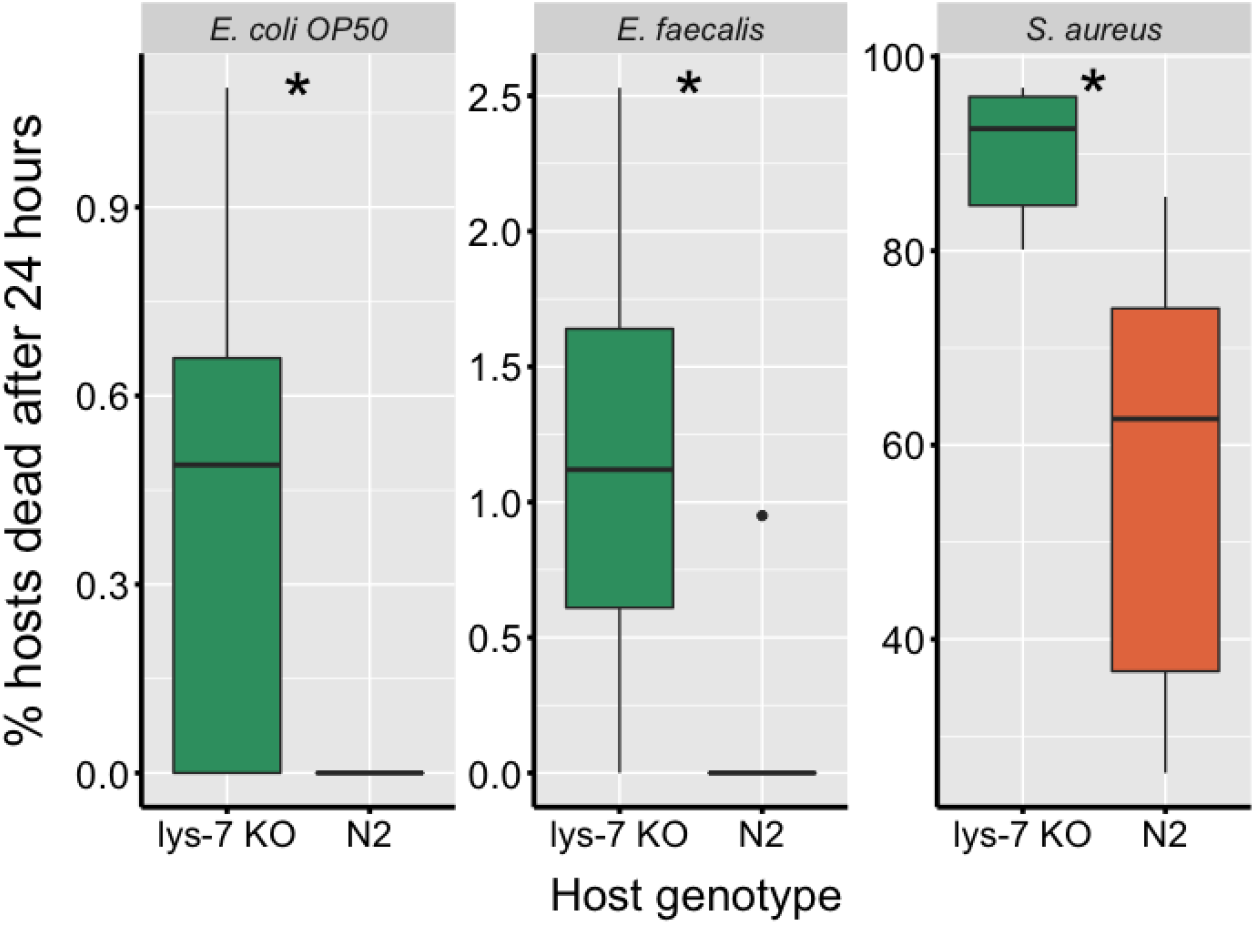
The effect of the *lys-7* gene on *C. elegans* mortality upon bacterial exposure. Percentage mortality of the wild-type N2 and *lys-7* knock-out mutant after exposure to pathogenic *S. aureus*, protective *E. faecalis*, and OP50 food. Each treatment was replicated five independent times. *P<0.05, **P<0.01, ***P<0.001.

## Discussion

Microbes can protect hosts from pathogens via altering the host’s immune response(4). They can also suppress pathogens in more direct ways that diminish the need for host-based immunity(1, 2). During the formation of a new symbiosis, these contrasting impacts of microbe-mediated protection on the immune system are important for predicting host vulnerability to infection should the interaction break down in future generations (2, 22). We investigated the effect of a bacterium with protective traits (*E. faecalis*)(25, 26) on the transcriptional response of a novel host (*C. elegans*) to infection by the pathogen *S. aureus*. We observed the upregulation of many immune/defence genes in the worm host such as those coding for the production of conserved antimicrobial proteins, including c-type lectins and lysozymes(36). This result reflects observations of other protective microbes found naturally (4, 37) or artificially (38) which modulate the host immune response.

Although other mechanisms of protection may still play a role in our system at the origin of the interaction, the transcriptomic evidence indicates that the immune system is a major factor. Here, *lys-7* was the most significantly upregulated lysozyme gene in protected hosts under attack by pathogens. Lysozymes are well conserved across the animal kingdom(30) and aid in resistance to *S. aureus* within *C. elegans*(31, 39). It was previously demonstrated in *C. elegans* that *S. aureus* drives the downregulation of *lys-7* expression and when expression is restored through chemical treatment, pathogen-induced host mortality significantly decreased along with infection load(39). Given *E. faecalis* is relatively more resistant to lysozyme activity(32, 33), we hypothesized that *lys-7* upregulation contributed to this bacterium’s protective effect. In support of this, we found that expression of this gene negatively affected *S. aureus* more than *E. faecalis*, providing a competitive edge to the protective microbe. Increased *E. faecalis* colonisation has previously been shown to relate to enhanced protection in our system (27, 40). These data are striking because the protective microbe was able to exploit host immunity in a novel species. This result may have emerged because *E. faecalis* is widely distributed among animal microbiota – as a protector (25), commensal and pathogen(41) – and lysozymes are conserved aspects of the animal immune response. Bacterial adaptations to these immune components in one host species might therefore be relevant in another. Indeed, the transfer of protective microbes to new hosts is more successful between host species that are more phylogenetically similar, i.e. their immune system is likely to be more similar(42).

Our results suggest that immune-mediated protection can occur in novel interactions, and could therefore be common at the beginning of a protective symbiosis. This finding has important implications for predictions on the evolution of host dependence and can be used to inform biocontrol design. Where protection involves the host immune system, it is predicted that hosts will continue to invest in their immunity against pathogens over evolutionary time(23). For biocontrol efforts, this is important to consider if the protective microbe should ever be lost from the host population, such as via imperfect transmission(43), altered environmental conditions (e.g. temperature change(43) or exposure to antibiotics (44)), or even the breakdown of protection (e.g. if the symbiont evolves to become pathogenic(2, 22)).

## Materials and Methods

### Nematode host and bacteria

*Caenorhabditis elegans* is a nematode that ingests microbes for nutrients (45–48) and is a well-established model for studying microbial colonisation and pathogenesis (37, 49–52). We used the simultaneous hermaphroditic N2 wild-type *C. elegans* strain from the Caenorhabditis Genetics Centre (CGC, University of Minnesota) along with a knock-out mutant for the *lys-7* gene that is outcrossed into the N2 genetic background (strain CB6738, CGC). We generated genetically homogenous lines by selfing a single hermaphrodite for 5 generations and froze them in 50% M9 solution and 50% liquid freezing solution in cryotubes at −80°C (50). We regularly resurrected populations throughout experimentation to prevent the accumulation of *de novo* mutations in host populations. Worms were maintained at 20°C on 9cm nematode growth medium (NGM) plates seeded with *Escherichia coli* OP50. *E. coli* OP50 is grown at 30°C shaking at 200rpm overnight in LB and 100μl of this is spread onto NGM plates and incubated overnight at 30°C(50). To ensure clean stocks and to synchronise the life stages of populations for experimentation, we treated worms with bleach (NaClO) and sodium hydroxide (NaOH) solution which kills everything except unhatched eggs (50).

We used *Staphylococcus aureus* strain MSSA 476 (GenBank: BX571857.1), an invasive community-acquired methicillin-susceptible isolate and *Enterococcus faecalis* strain OG1RF (GenBank: CP002621.1), both isolated from humans. We grew each species from a single colony overnight in 6ml Todd Hewitt Broth (THB) shaking at 200rpm at 30°C. Bacteria were frozen in a 1:1 ratio of sample to 50% glycerol solution in cryotubes at −80°C.

### Experimental set-up and RNA extraction

To examine host gene expression, we first exposed womrs to the protective microbe and pathogen or the pathogen alone, followed by RNA extraction. Sterile and age-synchronised eggs were collected using the bleach-sodium hydroxide solution. These eggs were kept in M9 buffer without food, shaking for ~8 hours at 88rpm and 20°C to arrest development at L1. Approximately 5,000 worms per replicate population were then transferred to 9cm NGM plates seeded with *E. coli* OP50 and placed at 20°C for two days. We chose this number to avoid overcrowding and starvation. We then grew both bacteria species from frozen culture overnight in 6ml THB in a shaking incubator at 30°C at 200rpm and diluted the optical density of *S. aureus* culture to be the same as the *E. faecalis* culture. We spread 120μl culture per species onto 9cm Tryptic Soy Broth (TSB) agar plates and incubated them overnight at 30°C. Where worms were to be exposed to both *E. faecalis* and *S. aureus*, we mixed 120μl of each culture together on the same TSB plate. We then removed worms from NGM plates, washed them in 50ml M9 buffer five times and placed approximately 2,000 young adults on the exposure plates at 25°C for 12 hours.

After 12 hours of exposure, we washed worms off each plate using M9 buffer within 10 minutes in an order determined by a random number generator. We chose 12 hours of exposure to avoid host mortality but provide sufficient time for *C. elegans* to respond to bacterial exposure and infection. We washed the worms in 10ml M9 buffer five times and put approximately 1,000 worms in 50μl into eppendorf tubes containing 1ml Trizol and vortexed for 20s. We then freeze-thawed the samples of worms three times using dry ice and a heat block (40°C) to break the worm cuticle and stored at −80°C. We extracted RNA using Zymo spin columns, following the manufacturer’s instructions with on-column DNA digestion using DNase I.

### RNA sequencing

We quantified the resulting RNA using the Qubit^®^ Fluorometer (Invitrogen) and all samples were diluted to the same final concentration. The Oxford Genomics Centre then performed library preparation and sequencing. The polyA signal was used to select the mRNA fraction from the RNA and this was converted to cDNA. Second strand cDNA synthesis incorporated dUTP and the cDNA was then end-repaired, A-tailed and adapter-ligated. Prior to amplification, samples underwent uridine digestion. The prepared libraries were size selected, multiplexed and checked for quality before paired-end sequencing using NovaSeq6000 with 150bp paired end reads.

We checked raw reads for quality using FastQC (0.11.5). Current (release 96) GTF and cDNA FASTA files were downloaded from the ensemble database for *C. elegans* (WBcel235 version of the *C. elegans* reference genome). We created a transcript index using Kallisto and the *C. elegans* WBcel235 cDNA FASTA file. We then performed pseudoalignment using Kallisto with 100 bootstraps and calculated transcript abundances. We then used the R package, Sleuth to perform statistical analyses (see ‘statistical analysis’).

### Host mortality

We tested whether the *lys-7* gene played a role in host mortality against *S. aureus* and *E. faecalis* independently and during co-colonisation. We grew age-synchronised eggs to young adult stage on 9cm NGM plates seeded with a lawn of *E. coli* OP50. Using the same protocols as describe above, we grew *S. aureus* and *E. faecalis in vitro* overnight, standardised the cultures to the same density, and made the exposure plates. We washed approximately 250 young adults of either CB6738 (*lys-7* knock-out) or N2 wild-type in 50ml M9 five times before exposing them at 25°C for 24h. The proportion of dead worms were then counted as a measure of host mortality.

### Bacterial colonisation

We exposed CB6738 (*lys-7* knock-out) and N2 wild-type worms to both *S. aureus* and *E. faecalis* together following the same protocol as above (see ‘Host mortality’). After 24 hours of exposure to the bacteria, we collected 7-10 live worms per exposure plate and washed them in 5ml of M9 five times under the microscope. To release the colonising bacteria, we placed the worms in 2ml screwcap tubes with 50μl M9 and 1.5mm Zirconium beads (Benchmark Scientific) and broke the cuticle by shaking the tubes at 320rpm for 45s. We plated serial dilutions onto Mannitol Salt Agar to isolate *S.aureus* and TSB with 100 μg/ml rifampicin (Sigma-Aldrich) to isolate *E. faecalis* and incubated the plates at 30°C overnight before counting the resulting colony-forming units (CFUs) per host.

### Statistical analysis

We performed differential expression analysis on the transcript abundance outputs from Kallisto using Sleuth in R v 3.2.0 (http://www.r-project.org/), using a likelihood ratio test of fitted models. The significance of treatment was determined by a q-value of <0.05 (p-value adjusted by means of the Benjamini-Hochberg false discovery date, FDR, correction for multiple comparisons). We performed a GO-term enrichment analysis on the significant differentially expressed genes using the g:Profiler online tool with the Benjamini-Hochberg FDR correction for multiple comparisons(53).

We used parametric tests for all data which met the required assumptions. We used the Shapiro test to detect whether data was normally distributed and F-tests to compare the variances of two samples from normal populations. We compared bacterial CFUs per host using two-sample t-tests. We used a binomial GLM to compare host mortality among host strains colonised by *E. faecalis* and OP50. To account for overdispersion, we used quasibinomial GLMs to compare host mortality among co-colonised host strains and host strains colonised by *S. aureus*. We checked GLM plots by eye for model quality and corrected p-values where multiple comparisons were made from the same dataset using the FDR (Benjamini & Hochberg) method.

## Data accessibility

Raw read transcriptional data will be deposited in the European Nucleotide Archive. The CSV files containing data for each figure will be uploaded to the Dryad repository.

## Acknowledgements

K.C.K. acknowledges funding from the Wellcome Trust (204826/Z/16/Z) and a European Research Council Starting Grant (COEVOPRO 802242).

## Author contributions

S.A.F. and K.C.K. conceived and designed the project; S.A.F. conducted the experiments and analysed the data; S.A.F. performed the bioinformatics analyses; S.A.F. and K.C.K. wrote the paper.

